# MarkerCapsule: Explainable Single Cell Typing using Capsule Networks

**DOI:** 10.1101/2020.09.22.307512

**Authors:** Sumanta Ray, Alexander Schönhuth

## Abstract

Many single cell typing methods require manual annotation which casts problems with respect to resolution of (sub-)types, manpower resources and bias towards existing human knowledge. The integration of heterogeneous data and biologically meaningful interpretation of results are further current key challenges. We introduce *MarkerCapsule*, which leverages the landmark advantages of capsule networks achieved in their original applications in single cell typing. Thereby, the small amount of labeled data required and the naturally arising, biologically meaningful interpretation of cell types in terms of characteristic gene activity patterns are exemplary strengths, beyond outperforming the state of the art in terms of basic typing accuracy. MarkerCapsule is available at: https://github.com/sumantaray/MarkerCapsule.

## Introduction

Single cell RNA sequencing (scRNA-seq) technologies provide unprecedented opportunities to capture high-quality gene expression snapshots in individual cells. Technological advances have recently enabled to process several thousands of cells per scRNA-seq experiment^1^. A fundamental step in single-cell analysis is to type the individual cells one analyzes. The most immediate approach is to cluster the population of cells under analysis into different groups. The groups to which individual cells belong then further determine the identity of the individual cells^2, 3^. This way of typing and annotating cells (reflecting *unsupervised learning* approaches) has been prevalent in identifying biologically coherent populations of cells in scRNA-seq data so far^4–7^.

However, the process of assigning biological meaning to cell clusters is both complicated and time-consuming, because it requires to inspect the identified cell clusters manually. Arguably, the procedure may even cancel the very advantages of single cell typing, because manual annotation requires to rely on prior knowledge, which had typically been gained by analyzing bulks of cells, and not single cells. Clearly, single cell experiments themselves constitute optimal resources for revealing and defining novel cell types^8^. With the revelation and availability of ever more cell sub-/types, cell states, possibly even at the level of cells transiting between types and states, manual inspection will be too tedious, inaccurate, or just impossible in terms of manpower resources. New methodology is required that allows to determine cell labels (sub-/types, states) in an automated fashion. *Supervised learning* based approaches^9–12^ address these points by being able to determine the identity of single cells although the characteristic molecular mechanisms have not yet been fully understood. This explains their recent gain in popularity^13^.

Tapping additional resources, such as protein expression data (e.g. sc-CITE-seq data^14^), methylation or chromatic accessibility data (the latter provided by e.g. sc-ATAC-seq^15^) and combining them with basic sc-RNA-seq data offers further advantages. When integrating additional data appropriately, one can expect both enhanced basic classification and the identification of cell (sub-)types that remain invisible on the RNA level alone. The driving methodical challenges are the heterogeneity of data sets, and the lack of general models for coherent integration^8^.

In this paper, we address the two challenges of establishing advanced supervised learning based methodology and the smooth integration of additional data in single cell typing. Beyond addressing these challenges in themselves, we also focus on their combination.

As for supervised learning, we employ capsule networks, as a most recent and advanced deep learning approach that has proven its superiority in other, prominent areas of application so far. Here, we demonstrate how to use capsule networks to equally great advantage also in single cell typing. We leverage the trade marks of capsule networks for typing cells at utmost accuracy, economic use of training data (Capsule Networks achieve competitive accuracy already on only half or even only a third of the data in comparison with state-of-the-art approaches), and biologically meaningful interpretation of results. Note that the reduced demand of training data helps to identify also types where cells suffer from a relative lack of coverage. Biological interpretation addresses the notorious complaint that high-performance deep learning is little explainable: here we can deliver explanations along with outstanding performance.

We also present novel technology that enables a coherent integration of additional data. We create data representations based on non-negative matrix factorization and variational autoencoders that support the coherent integration of data from additional experimental resources, such as sc-ATAC-seq, sc-CITE-seq, or sc-bisulfite sequencing, and which is generally applicable. Experiments point out that these representations enhance the performance in classification even further. That is, we demonstrate that mastering the methodical challenges of the integration of additional data indeed means an important step up in classifying single cells, as was anticipated in earlier studies.

### Summary of Contributions

In this work, we provide the following novelties:

1. We provide the first capsule network based approach (MarkerCapsule) that successfully deals with sequencing data in general, and single cell sequencing data in particular. Note that earlier Capsule Network based biological applications have only addressed protein structure prediction^16^, the prediction of secretory proteins in saliva based on non-sequencing based proteomics data raised in 2007/8^17^ and in network and disease biology^18^ (where their possible advantages for processing heterogeneous, multi-omics data were pointed out).
2. Although studying the integration of additional data has gained considerable momentum recently^8^, our approach is the first *supervised* learning approach that is explicitly designed to integrate data from multiple single cell analysis techniques. While the state of the art in supervised learning based single cell typing^9–12^ is able to process integrated data when suitable data representations are provided, our approach is the first one to explicitly provide such representations. For earlier *unsupervised* approaches that deal with integration, see^5, 19, 20^ for combined measurements on transcription, chromatin accessibility and/or genome methylation, and^21, 22^ for combining scDNA and scRNA sequencing.
3. Our capsule network based approach outperforms all supervised learning methods that are state of the art in single cell type classification on integrated data. This confirms that the theoretical achievement (2) yields relevant practical advantages as well.
4. We demonstrate that MarkerCapsule requires (substantially) less training data than prior approaches for achieving optimal performance. This enables to identify sparsely covered cell types, without incurring losses in prediction accuracy.
5. We demonstrate that the primary capsules of the MarkerCapsule network have a clear and intuitive interpretation: we show that primary capsules reflect marker genes for cell types where marker genes are known. If marker genes are not known for a cell type, genes captured by primary capsules that are important for predicting such types are likely to suggest plausible marker genes.

## Results

### Overview

In the following, we present our new method, *MarkerCapsule*, and explain its overall *workflow*. MarkerCapsule’s foundation are *capsule networks*, which reflects most recent breakthrough technology in deep learning. We proceed by outlining the mechanisms and intuition of capsule networks, which had their greatest achievements in image classification so far. We then continue to describe the methods that we use for the *integration of additional data*, the basis of which are non-negative matrix factorization and variational autoencoders.

Subsequently, we demonstrate in *experiments* that *(i)* MarkerCapsule outperforms the state-of-the-art in basic scRNA-seq based single cell typing, *(ii)* the integration of additional data enhances the classification of MarkerCapsule further, *(iii)* in MarkerCapsule, the primary capsules, as fundamental building blocks of Capsule Networks, refer to sets of marker genes for all types of cells where marker genes are known (which suggests that primary capsules of MarkerCapsule capture marker genes in general), and *(iv)* that MarkerCapsule requires substantially less training data than other approaches to achieve optimal performance in prediction.

### MarkerCapsule: Workflow

See Figure 1 for the workflow of our analysis pipeline. We describe all important steps in the paragraphs of this subsection.

**Figure 1.**
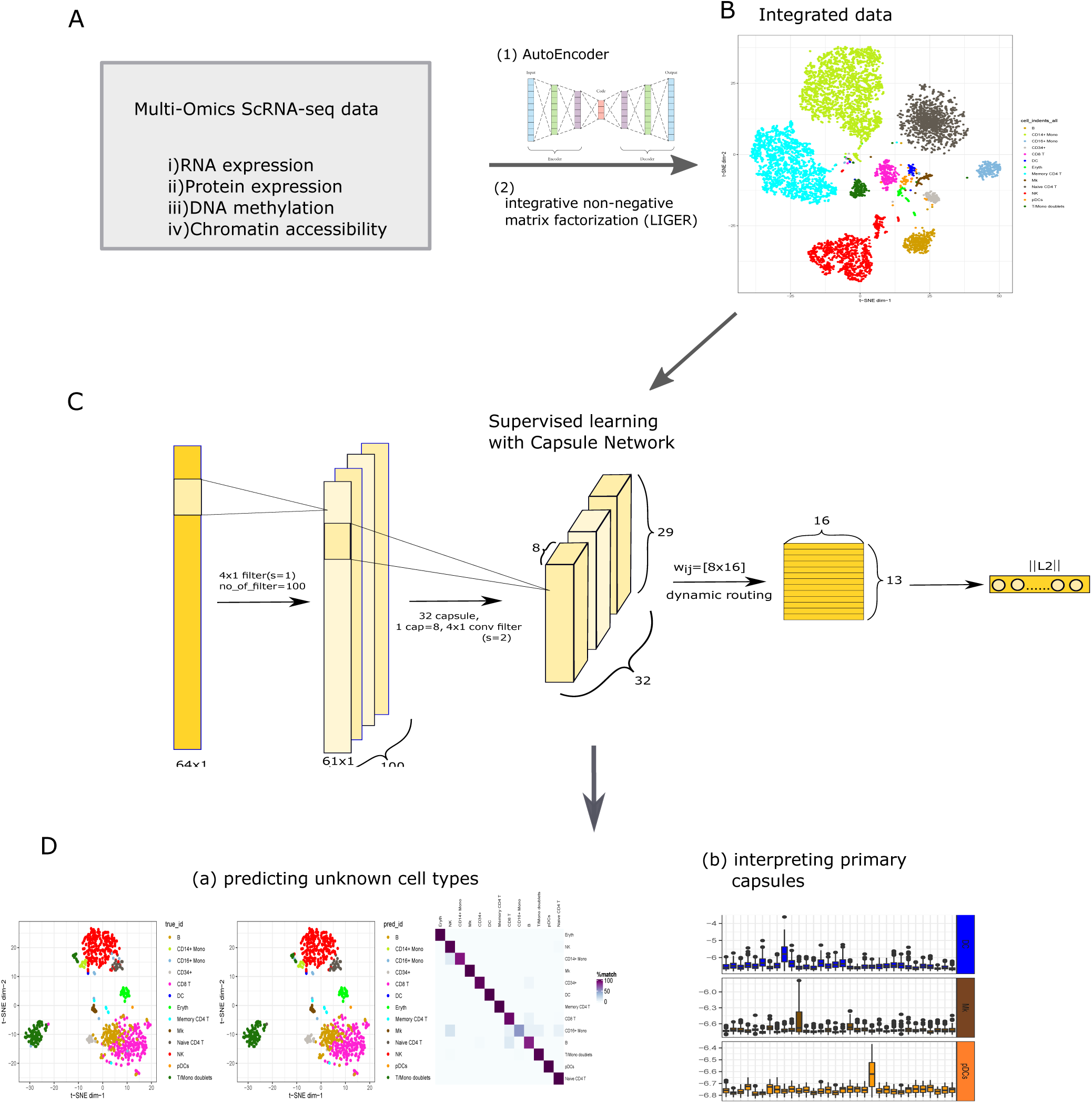
Workflow of the MarkerCapsule framework.

#### Input and integration of heterogeneous data

See Panel A and B of Figure 1. The basic input for MarkerCapsule are scRNA-seq data that measure gene expression. If data referring to different experimental techniques on the same single cells are available, integration of the data prior to running MarkerCapsule is necessary. The challenges are the heterogeneity of the data sets, and the lack of general models that address how to join heterogeneous data in a meaningful way. Here we use LIGER^23^, an integrative non-negative matrix factorization (NMF) based approach, on the one hand, and a variational autoencoder (VA) based technique, on the other hand, to integrate additional data (DNA methylation, protein expression, chromatin accessibility) with the expression data; see ‘Datasets’ in Methods for a detailed description of the four different integrated data sets we used.

#### Training and optimizing MarkerCapsule

See C in Figure 1. As usual, we divided the full data sets into training, validation and test data. For exploration and adjustment of hyperparameters and optimizing the underlying capsule network architecture, we evaluated MarkerCapsule in cross-validation runs. The resulting capsule network that underlies MarkerCapsule consists of an ordinary convolutional layer, followed by a primary capsule and an output capsule layer (the connection between the latter two of which encompasses a dynamic routing procedure). Output capsules refer to the individual cell types that one can predict. See Methods for details.

#### Predicting cell types with MarkerCapsule

See D (a) in Figure 1. For the evaluation of the performance of MarkerCapsule, we consider the test data from the basic scRNA sequencing gene expression data sets and the data sets that integrate scRNA-seq gene expression data with data that refer to protein expression, DNA methylation, chromatin accessibility (scATAC-seq) or additional, complementary scRNA-seq gene expression data. After final training, the test data is provided as input to MarkerCapsule. MarkerCapsule’s performance is evaluated in terms of the accuracy with respect to predicting cell types of the test data correctly.

#### Interpreting the primary capsules

We demonstrate that the primary capsules of MarkerCapsule allow for a biologically meaningful interpretation of the results: we experienced that the primary capsules correspond to marker genes for cell types for which marker genes are available. For cell types for which such marker genes are lacking, the genes that contribute the most to the relevant primary capsules may correspond to novel marker genes. Note that the lack of a meaningful interpretation of results is usually considered a crucial drawback of deep learning. Here, we overcome this issue by associating MarkerCapsule’s fundamental network building blocks with genes that in their combination act as cell type specific marker genes.

### Capsule Networks in Single Cell Analysis

*Capsule Networks (CapsNets)* are a deep neural network architecture, originally presented in 2017/18 and applied in digit recognition and image classification (by the pioneers of deep learning themselves^24^). CapsNets established great advantages over the prior, usually convolutional network based state of the art, resolving fundamental, year-long standing methodical issues. The predominant achievements of CapsNets were, *first*, to achieve superior performance in classifying/recognizing overlapping or distorted images, *second*, to allow for complex, while still human-mind-friendly interpretation of their primary building blocks (the *capsules*, as particularly arranged groups of ordinary neurons, which in digit recognition, for example, referred to stroke thickness, gray scale, and so on), and *third*, to require substantially less annotated (training) data for achieving outstanding performance.

Note that all of these 3 points refer to major points of critics about deep learning. In that, CapsNets overcame the major drawbacks that were commonly attributed to deep learning. In single cell typing, pixels correspond to gene expression values. Following this analogy:

- Overlapping image elements correspond to (groups of) genes that support multiple, possibly interacting or interfering cellular pathways. The corresponding intuition is that Capsule Networks are able to identify cell types whose defining elements are such interacting pathways (while other methods tend to stumble across such interference).
- Primary capsules correspond to groups of cooperating genes in single cell typing. We demonstrate that the corresponding genes correspond to marker genes, in all cases where such marker genes were available in the literature (which justifies the name “MarkerCapsule”). This suggests that primary capsules may capture suitable marker genes for cell types for which marker genes have remained unknown.
- Last, CapsNets required substantially less training data for accurately predicting cell types. So, CapsNets can accurately identify cell types that are only sparsely covered by data, while other methods tend to get confused with such types.

See Methods for further facts and information about Capsule Networks.

### Integrating Additional Data

For the integration of scRNA-seq data with data from other (sequencing based) single cell experimental resources, we have utilized two methods. *First*, LIGER^23^, as a *non-negative matrix factorization (NMF)* based strategy originally used in clustering single cells based on multiple data sets and, *second*, a *variational autoencoder (VA)* based strategy originally designed for cancer data integration^25^ which we developed further for the particular purposes here.

Features in the original data sheets refer to read counts reflecting gene expression values for scRNA-seq data on the one hand, and read counts reflecting degree of methylation or protein expression, and so on, for the additional experimental data, on the other hand (see ‘Data Sets’ in Methods for detailed descriptions of the data sets under consideration in our experiments). Features computed by applying NMF or VA to several data sets together reflect combinations of features that are optimized with respect to correctly differentiating between the different cell types. To represent the original data in terms of these new features, all data is processed by NMF or VA. The resulting data representation is then passed through the capsule networks.

Note that NMF, while traditionally being very powerful, requires to process all available data in one run so as to obtain integrated representations that apply for both labeled and unlabeled data. Subsequently, one trains the capsule network on the integrated, labeled training data, and predicts cell types by running the trained network on the integrated unlabeled test data. In the following, our experiments reflect this procedure. Online NMF and/or supervised NMF are active areas of research^26, 27^, which promises that soon integrated NMF based representations can be inferred from labeled data alone and subsequently applied to unlabeled data, which avoids re-training runs. VA, unlike NMF, reflect generative models and therefore can be inferred only from the labeled training data, and only subsequently applied to also unlabeled data. This evidently optimally caters to supervised learning protocols. For utmost consistency in terms of comparability of results, however, we apply the protocol used for NMF also for VA.

In summary, the advantage of integrated features is to exploit effects that become evident only after combining the heterogeneous data sets. This unmasking of additional effects explains the increase in accuracy in predicting cell types.

### State-Of-The-Art Methods

We compare MarkerCapsule with the current state-of-the-art in supervised learning based single cell typing. We recall that supervised learning reflects an advantage over unsupervised learning (clustering) because it enables to automatize the annotation step, which explains why we focus on methods that do not require a manual annotation step. Note that all state of the art methods are able to take integrated data as input. As a side remark, it may speak for the new data representations that all prior approaches get boosted by them, although none of these approaches originally intended to deal with integrated data. In the following, we consider **(1)** *Scpred*^12^, which combines an unbiased feature selection method with standard machine-learning classification, where dimensionality reduction and orthogonalization of gene expression values prove advantageous to accurately predict cell types **(2)** *AcTINN*^11^ (Automated Cell Type Identification using Neural Networks), which utilizes a neural network model with three hidden layers, trained and used for prediction the usual way **(3)** *CHETAH*^28^, which builds a hierarchical classification tree from the reference (training) dataset, and classifies unknown samples by computing the correlation between genes that discriminate the test cell from the reference dataset and, finally, a basic CNN that serves the purposes of a generic deep learning background model. **(4)** Note that *Garnett*^9^, as another state-of-the-art approach requires to supply a cell type marker file, along with the usual training data, which does not suit the general setting of supervised learning based single cell typing. **(5)** As for a CNN baseline model, we choose two convolution layers followed by two dense layers, which matches the Capsule Networks in terms of numbers of layers of capsules. For the first layer, we use 4×1 filter with stride 1, and for the second layer, we choose the same filter size with a stride of 2.

See Methods for the corresponding choices of parameters, which generally reflect default settings.

### Advantages of MarkerCapsule in single cell typing over state of the art

We compared MarkerCapsule with the state of the art methods (Scpred, AcTINN, CHETAH; see above) and the baseline CNN model on the basic scRNA-seq expression data. All such basic data sets were taken from the four data sets we are considering; see ‘Data Sets’ in Methods for a detailed description of the data and numbers of samples. Note that in this subsection, we do not integrate the basic scRNA-seq data with the additional data that are provided as parts of the four data sets. The purpose is to be able to evaluate how accuracy in prediction compares between using only basic scRNA-seq data and integrated, heterogeneous single cell sequencing data.

Each basic scRNA-seq data set is divided into training, validation and test data at a ratio of 8:1:1. Cell type prediction performance is evaluated by determining the average test accuracy and the corresponding standard errors over 100 runs for each of the competing methods. For exploring how the different approaches react to reducing the training data, data was subsampled at rates ranging from 10% to 100% in steps of 10% prior to training, where random subsamples were drawn for each of the 100 runs independently.

See Figure 2 for the corresponding results. MarkerCapsule largely dominates the other methods across the varying sizes of training samples. The only method that puts up real competition is ACTINN, which nearly equalizes the accuracy in prediction of MarkerCapsule but only when full data sets are provided for training. Still, MarkerCapsule outperforms also ACTINN substantially when smaller data sets are used for training.

**Figure 2.**
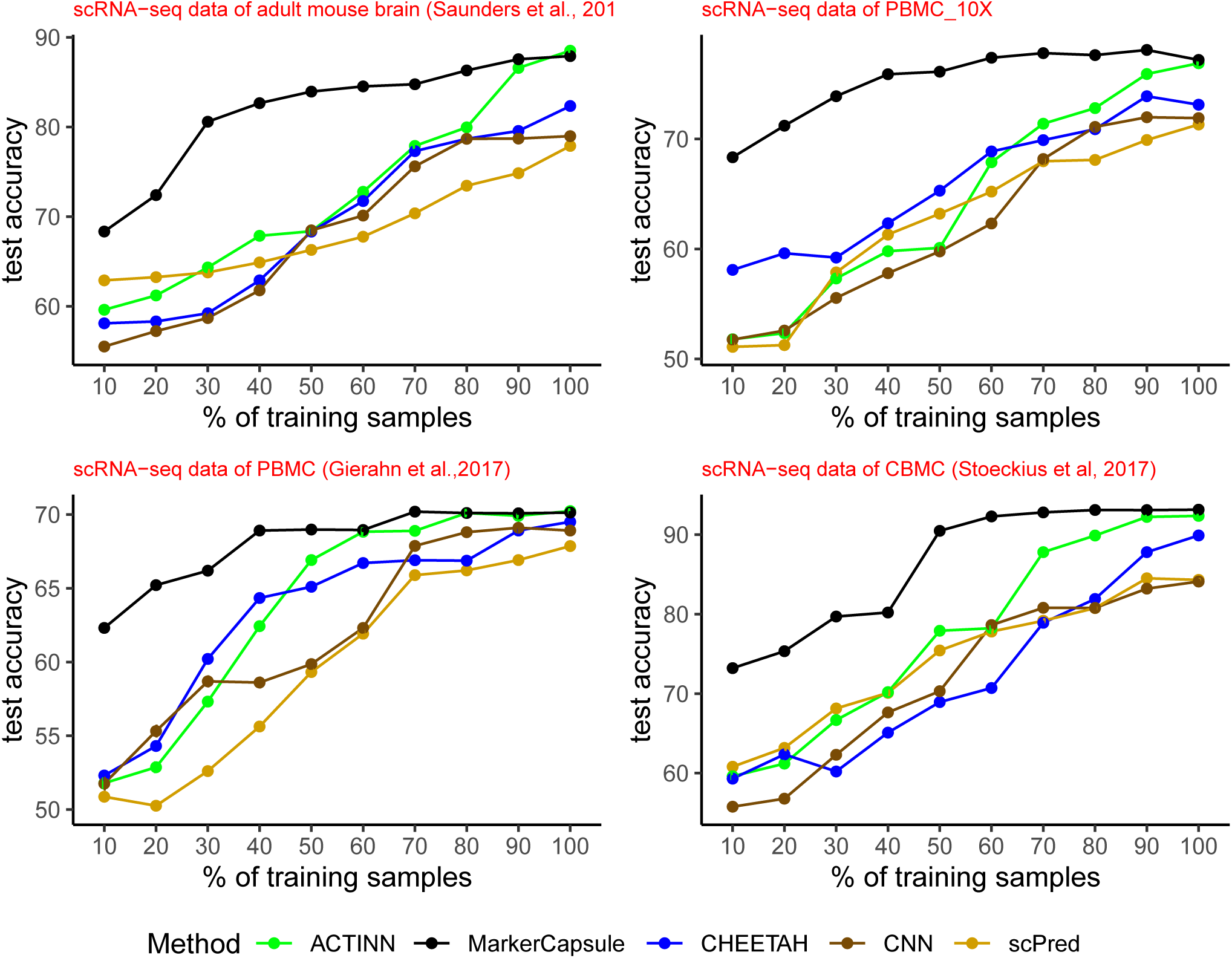
Performance of different methods on basic scRNA-seq datasets. Average test accuracy is reported over 100 runs for each of the competing methods

### MarkerCapsule enables to exploit heterogeneous, integrated data

We further evaluated the effects of integrating additional data using NMF and VA based strategies, as discussed above. Therefore, we made use of the full four data sets, as described in ‘Data Sets’ in Methods. The different data sets integrate protein expression data (CITE-seq), methylation data, scATAC-seq data or additional scRNA-seq data that was sequenced independently on a different platform.

As for basic scRNA-seq data, we divided also each integrated data set into training, validation and test data, again at a ratio of 8:1:1. In each run, we computed integrated data representations only on the training data, and we proceeded by using these representations also for the validation and the test data, reflecting a realistic scenario in practice.

Again, we vary the size of the training samples by subsampling them from 20% to 100% at steps of 10%. We present test accuracy, as the average of test accuracy across 100 different runs for each method.

Table 1 shows the corresponding mean accuracy values, together with their standard deviation, with respect to the individual accuracy values across those 100 runs. First, it becomes evident that the accuracy of MarkerCapsule relative to determining the correct cell type is improved by about 3-4% on all data sets relative to the results obtained when running MarkerCapsule on only the basic scRNA-seq data. This provides clear evidence of the advantages of the exploitation of integrated data over using only basic scRNA-seq data.

**Table 1.** Table showing the performance of the four competing methods in four integrated datasets trained with different percentage of training samples. Maximum values are highlighted in bold font.

| Test accuracy of models trained on different % of training sample |  |  |  |  |  |  |  |  |  |  |
| --- | --- | --- | --- | --- | --- | --- | --- | --- | --- | --- |
| Method | Dataset-1: CITE-seq |  |  |  |  | Dataset-2: scRNA-seq + snATAC-seq |  |  |  |  |
|  | 20% | 40% | 60% | 80% | 100% | 20% | 40% | 60% | 80% | 100% |
| scPred | 63.81±0.36 | 70.13±0.29 | 78.92±0.20 | 82.81±0.09 | 89.26±0.03 | 51.25±0.21 | 62.32±0.20 | 65.46±0.21 | 68.71±0.12 | 70.41±0.10 |
| ACTINN | 61.29±0.33 | 70.38±0.25 | 79.80±0.19 | 91.29±0.11 | 95.03±0.10 | 52.14±0.35 | 63.42±0.31 | 67.81±0.22 | 73.85±0.20 | 76.71±0.19 |
| CHETAH | 62.37±0.29 | 65.89±0.11 | 72.71±0.15 | 83.91±0.10 | 90.34±0.05 | 60.91±0.30 | 62.34±0.25 | 67.63±0.21 | 71.29±0.10 | 75.61±0.08 |
| MarkerCapsule | **69.81±0.10** | **80.23±0.03** | **94.53±0.01** | **95.58±0.006** | **95.05±0.001** | **69.31±0.03** | **70.34±0.03** | **76.85±0.02** | **77.12±0.02** | **77.39±0.02** |
| Method | Dataset-3: scRNA-seq (10X + seq-well) |  |  |  |  | Dataset-4 (scRNA-seq + DNA-Methylation) |  |  |  |  |
|  | 20% | 40% | 60% | 80% | 100% | 20% | 40% | 60% | 80% | 100% |
| scPred | 50.29±0.31 | 58.91±0.30 | 62.10±0.25 | 66.57±0.19 | 67.86±0.05 | 65.87±0.27 | 66.48±0.23 | 68.25±0.20 | 73.03±0.10 | 77.81±0.05 |
| ACTINN | 50.91±0.49 | 62.30±0.43 | 64.51±0.35 | **68.90±0.19** | **70.31±0.09** | 63.21±0.31 | 68.91±0.29 | 74.56±0.19 | 81.29±0.19 | **89.32±0.10** |
| CHETAH | 54.32±0.22 | 60.38±0.20 | 62.19±0.10 | 67.35±0.11 | 69.82±0.07 | 61.37±0.28 | 63.45±0.22 | 72.71±0.29 | 81.29±0.10 | 86.89±0.07 |
| MarkerCapsule | **65.14±0.09** | **66.27±0.05** | **66.31±0.05** | 68.81±0.01 | 69.72±0.01 | **74.50±0.10** | **84.53±0.10** | **87.81±0.05** | **88.10±0.01** | 88.92±0.01 |

**Table 2.**
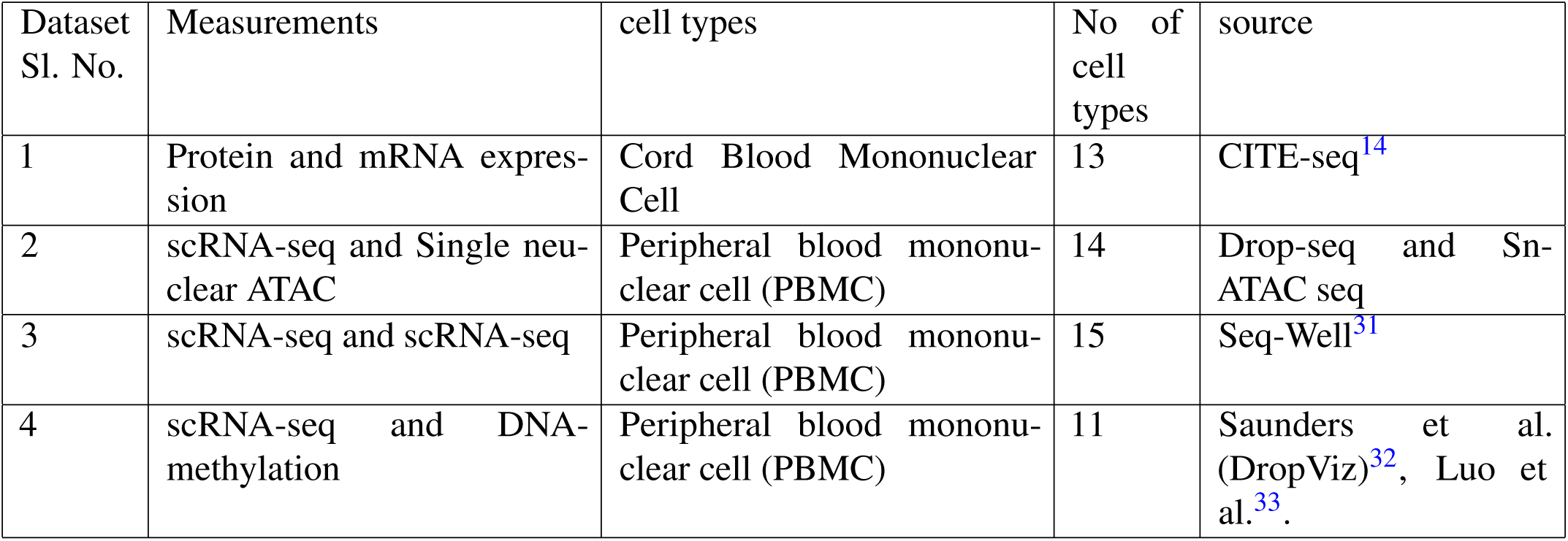
Description of data sets used in experiments

Also on integrated data, the only method that is able to compete with MarkerCapsule is ACTINN. However, in all but one case, ACTINN incurs substantial losses already when being confronted with only 80% of the original data (and incurring further losses when further decreasing the size of the training data samples). Moreover, MarkerCapsule is (substantially) more stable with respect to the different runs, documented by the fact that ACTINN’s standard deviation is approximately 10 to 100 times larger than that of MarkerCapsule. In summary, MarkerCapsule proves to be the approach that is most favorable in practice, as the one achieving optimal performance in all settings, and having substantial advantages with respect to training data resources required and stability of training runs relative to variations/fluctuations in the data provided for training.

Further, a t-SNE based visualization of original data and samples of the test data of Dataset-4 (integrated with CITE-seq, see Methods) as predicted by MarkerCapsule is shown in Figure 3, Panel A and B. Percentages of samples with correctly predicted labels are further shown in the heatmap of Figure 3, Panel C. The visualization documents that MarkerCapsule’s predictions (Panel B and C) become confused only in cases for which the data provides little evidence for distinction (Panel A).

**Figure 3.**
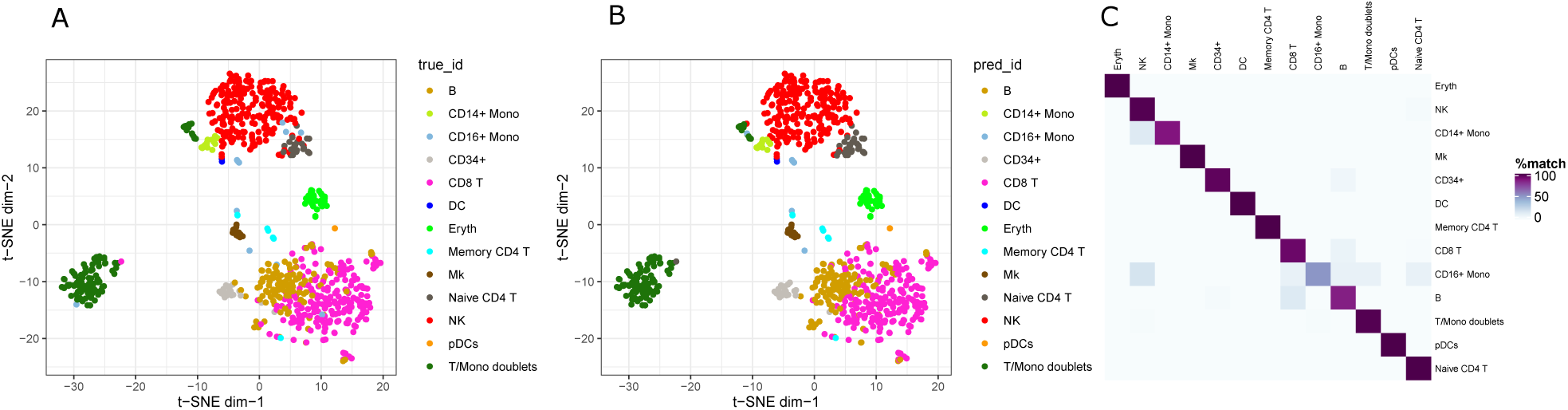
Figure shows performance of the proposed model in a random test sample of CITE-seq data. Panel-A-B represents the t-SNE visualization of the test samples with the original label and predicted cell types for integrated dataset-4. Pane-C shows the heatmap of percentage of matching between the original and predicted labels.

### Capsule Networks versus Convolutional Neural Networks

For further understanding the advantages, and also the possible drawbacks of capsule networks in comparison with ordinary convolutional neural networks, we conducted further experiments that focused on exploring the corresponding differences.

### Training and Validation

Panel A in Figure 5 displays the training and validation accuracy of the CNN, as described above ((5) in ‘State of the Art Methods’), on the one hand, and MarkerCapsule on the other hand, when running them on the four integrated data sets.

**Figure 4.**
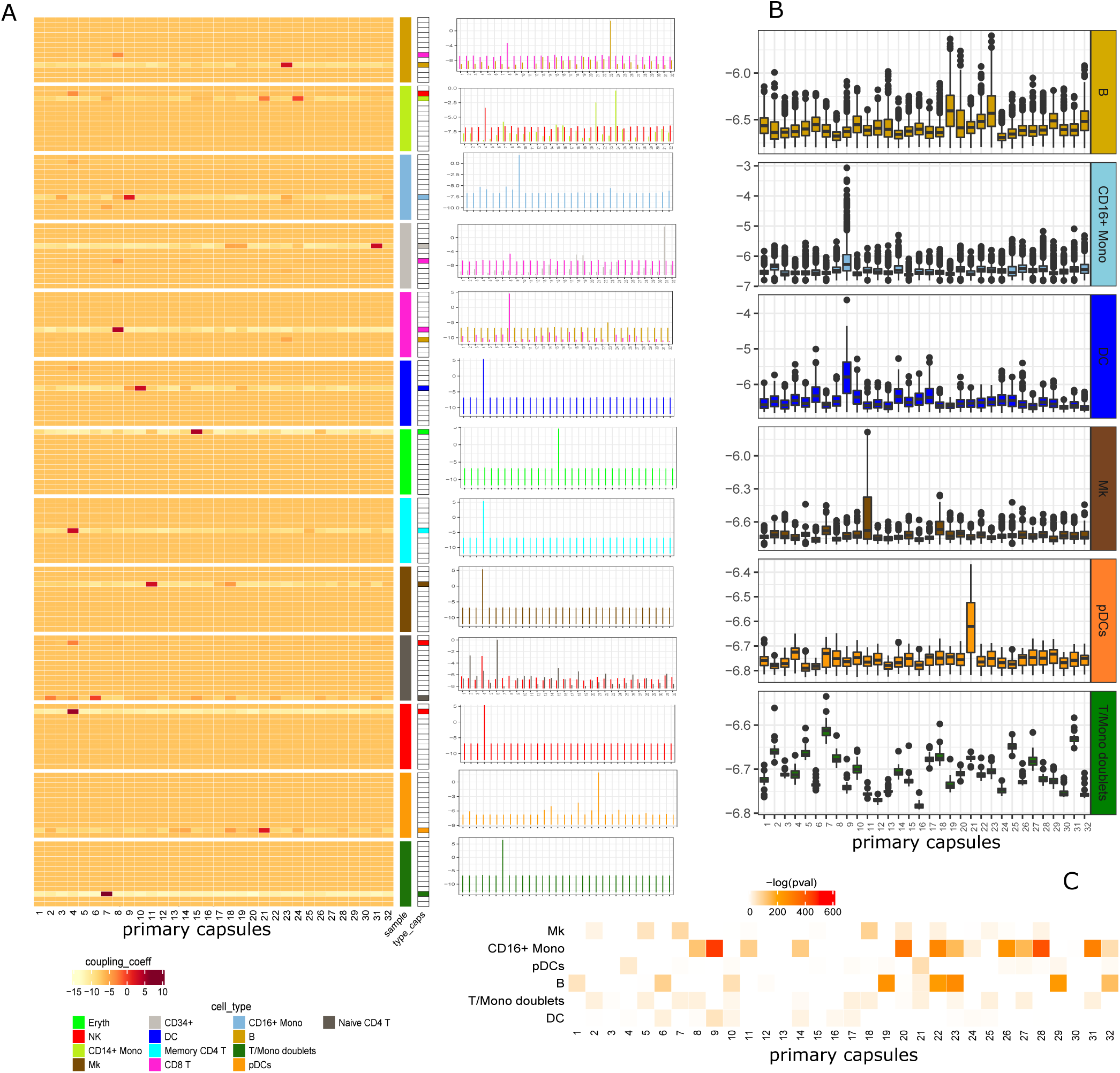
Figure shows the interpretation of capsules. Panel-A shows heatmap of average values of coupling coefficients for 100 different samples with distinct cell types. For each sample we have 13×32 dimension matrix of coupling coefficients and each matrix is visualize as a heatmap. In the right side of the plot the corresponding signal of the primary capsules are shown in spike plot. Panel-B shows boxplots of coupling coefficients for 6 cell types by taking only the expression of marker genes.

**Figure 5.**
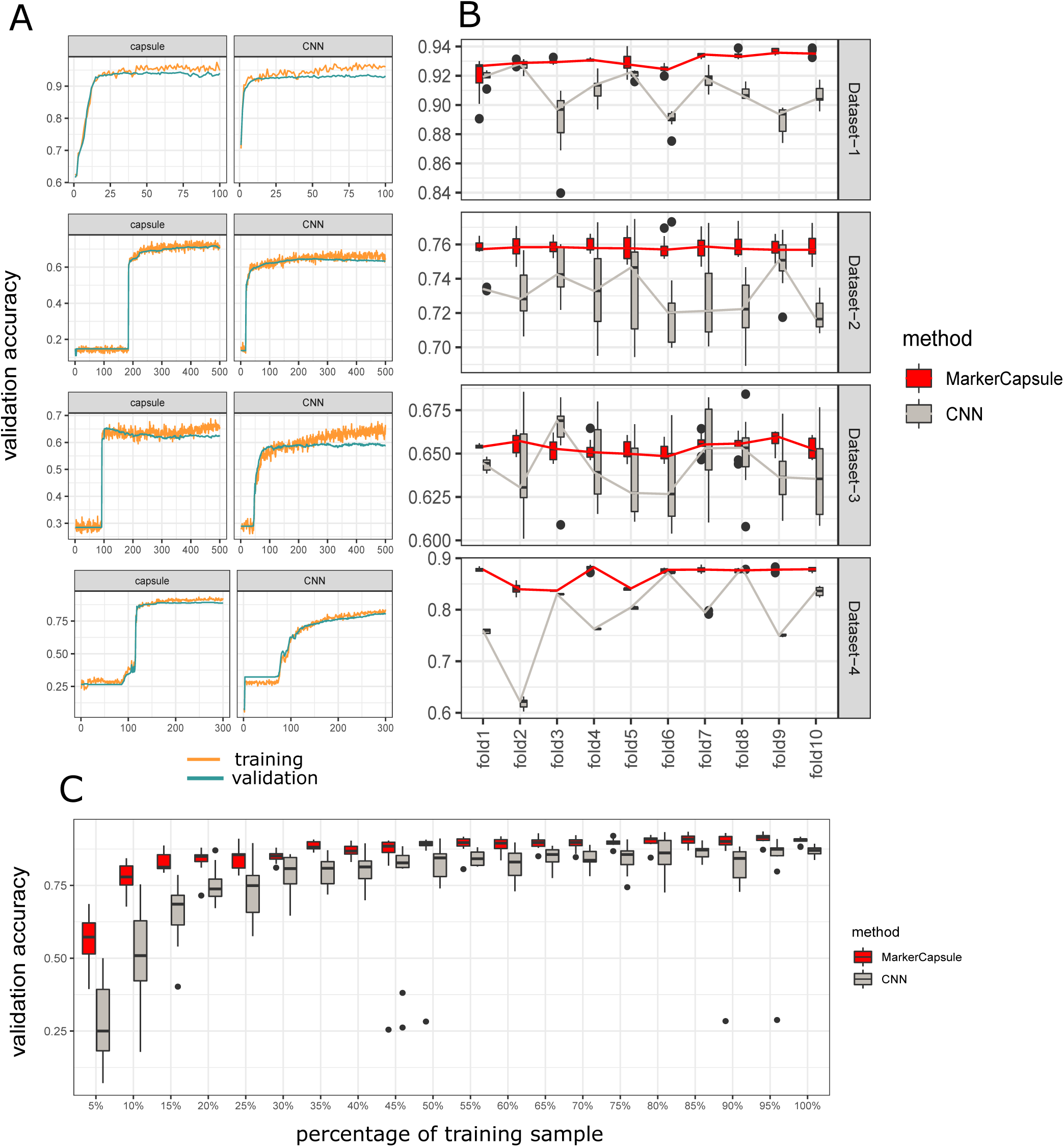
Performance comparison between MarkerCapsule and CNN model. Panel-A shows training and validation accuracy of both model in all four integrated datasets. Panel-B shows consistency of validation accuracy in 10-fold cross validation experiment. Panel-C represents validation accuracy of both models trained with different percentage of training samples.

The first observation is that MarkerCapsule consistently achieves superior accuracy in comparison with the CNN, which establishes an obvious and fundamental advantage.

A second observation is that MarkerCapsule can require substantially more time for training, as measured in terms of epochs needed until operative performance in validation accuracy is achieved. While the trend can be observed on all data sets, this is particularly striking on Dataset 2. This disadvantage reflects a traditional drawback of Capsule Networks: they required more time for training in their original application as well. The corresponding reasons are well known and understood. It is important to realize that one can observe no particular disadvantages in this respect when applying Capsule Networks to single cell typing.

An additional advantage of MarkerCapsule over CNN’s is the stability of its performance, not only with respect to varying training and validation data (as established through 10-fold cross validation), but also with respect to variations across epochs (as basic training data units). The box plots in Panel B, Figure 5 provide the respective evidence: on the y-axis validation accuracy is plotted, which varies across the different validation folds plotted along the x-axis. For each fold, the box indicates the mean and the standard deviation of the validation accuracy achieved for the respective fold, across the last ten training epochs. It is obvious that the mean accuracy of MarkerCapsule (across both folds and epochs) is clearly greater than that of the CNN’s. In addition the standard deviation across folds is substantially smaller (indicated by the red line being flatter than the gray line). Within each fold, the standard deviation is substantially smaller as well, indicated by the smaller size of the boxes.

In summary, MarkerCapsule proves to be both significantly more accurate and stable in comparison with CNN’s.

### Learning from little data

To understand how performance relates with the amount of training data provided, we evaluated the performance of both MarkerCapsule and the CNN relative to varying the size of the training data set in a setting that addresses this more explicitly (in comparison to the experiments from above). To this end, we split the data into training, validation and test data at a ratio of 8:1:1, as before. To implement cross-validation we varied training and validation data (while keeping the test data fixed). In each cross-validation fold, we evaluate both MarkerCapsule and the CNN while varying the size of the training data used for training. We start at 5% of the training data, and upon completion of a training run (where completion corresponds to 150 epochs for each training run, for the purposes of a consistent evaluation across folds and sizes, where 150 epochs proved to be sufficient in the experiments from above, not only for the CNN, but also for MarkerCapsule), we add another 5% of the training data for the subsequent training run, until full usage of training data in that cross-validation fold. For each size (5%, 10%, …, 95%, 100%), we collect the resulting validation accuracy across the cross-validation folds, and display mean and standard deviation of the fold-specific values in terms of box plots, see Panel C in Figure 5 (grey: CNN; red: MarkerCapsule).

It becomes evident that MarkerCapsule clearly outperforms the CNN, both in terms of mean validation accuracy and standard deviation. In other words, MarkerCapsule proves to be superior both in terms of basic performance and in terms of training stability. Note that both criteria—superior performance on small training data and improved stability—reflect characteristic advantages of Capsule Networks. The reason is that they depend on variations/fluctuations of the training data provided to a much smaller degree (as further explained by the “Viewpoint Invariance Property” that applies for capsule networks, see above and Methods). As for particular examples, consider that the performance of MarkerCapsule achieved on 10% of the training data is rivaled by the CNN only at 30% of the training data, or that the CNN requires the full data set to achieve the performance achieved by MarkerCapsule using only 35% of the training data. In general, MarkerCapsule requires only one third of the training data in comparison to standard CNN’s for achieving superior performance. Also these results reproduce characteristic advantages of Capsule Networks, as documented in various earlier applications.

In summary, the results document that MarkerCapsule leverages the landmark advantages of Capsule Networks, as achieved over CNN’s in earlier applications, also for single cell typing.

### Interpretation of primary capsules

#### Features learned by primary capsules

To understand whether the primary capsules of MarkerCapsule have the potential to capture effects that allow for an intuitive and biologically meaningful interpretation, we studied the coefficients attached to the edges that connect the primary with the output capsules (“type capsules”), commonly referred to as “coupling coefficients” (see Methods). Therefore, for each cell type, we randomly select 100 samples of that type from the test set of Dataset-1 (CITE-seq); if a cell type has less than 100 test samples, we consider all of them. We then run the previously trained ‘MarkerCapsule’ model on each of the selected cell samples.

We save the values for the coupling coefficients for each of the 100 samples (less if the test data for one type is smaller, see above; in the following, for sake of simplicity, we refer to the 100 samples for all the cell types). Because MarkerCapsule connects 32 primary capsules with 13 output (type) capsules, we obtain 13×32 coupling coefficients for one particular test cell sample, which amounts to 100×13 ×32 coupling coefficients overall, for one cell type overall. We compute the average values for the coupling coefficient across the 100 samples, which results in 13×32 averaged coupling coefficients for each of the 13 cell types, yielding 13×13×32 averaged coefficients overall. We collect these averaged coefficients into 13 matrices of size 13 ×32.

See Figure 4, Panel A for the corresponding 13 matrices (for 13 cell type) of size 13×32. For the sake of more distinct visualization, the probabilities *p* corresponding to the coupling coefficients are converted using the function log(*p/*1 −*p*)

In Figure 4 one observes that for each cell type at least one primary capsule is clearly activated. For example, primary capsule 15 is activated for cell type ‘Eryth’, while primary capsule 4 is activated for cell type ‘NK’. Some types have more than one primary capsule activated: for cell type ‘CD14 ±’ for example both primary capsules 21 and 24 are “firing”. At any rate, for each type, few primary capsules can be clearly distinguished that correspond to the type by their level of activation.

While the majority of types shows a clear activation of the correct type capsule, some type capsules are activated for two types. For example: for ‘CD8 T’, both ‘CD8 T’ and ‘B-cell’ capsules are active (where ‘CD8 T’ is more active than ‘B’), and for ‘CD34+’, both ‘CD34+’ and ‘CD8 T’ type capsules have been activated. However, in all such cases, the activation of the correct type capsule dominates the alternative type capsule such that the correct prediction of the type is still possible.

To further understand these ambiguous cases, we study the corresponding signals of the primary capsules that contribute to the cases, and plot the values of the coupling coefficients as barplots, shown at the right of the heatmaps in Panel A of Figure 4. For the cases where more than one type capsule is activated we observe that the primary capsule that correlates the strongest with the correct type fires the most, which explains why a clear distinction is still possible. For example, in the ‘CD8 T’ sample, where the ‘B’ cell type capsule is also activated, the signal of the primary capsule that correlates the strongest with the ‘B’ cell type-capsule (primary capsule 23) is much lower than the signal of the primary capsule that correlates the strongest with the ‘CD8 T’ type capsule itself (primary capsule 8).

### Relation of primary capsules with marker gene expression

We further investigated the hypothesis that the genes that contribute the most to primary capsules (reflected by input neurons that fire the most for primary capsules) represent marker genes for the types that the primary capsules predominantly support. To this end, we downloaded marker genes for several different cell types from the CellMarker database^29^. This way, we were able to extract marker genes that matched input gene sets to a sufficient degree for six cell types, namely ‘B’, CD16+ Mono’, ‘DC’, ‘Mk’ ‘PDcs’ and ‘T/Mono doublets’. For the remaining cell types the input gene set proved too small to match collections of marker genes to allow for a statistically reasonable investigation.

For each of the six cell types equipped with marker genes, we randomly selected 100 samples from the test set and applied the MarkerCapsule network trained earlier on these samples. The resulting coupling coefficients between primary capsules and the corresponding type capsules are plotted in Figure 4 Panel B. For each cell type in Panel B of Figure 4, one box captures the 100 coupling coefficients attached to the edge between the primary capsule and the type capsule, displaying the mean and the variance of the coupling coefficients across the 100 samples. For example, the first sub-panel, referring to cell type ‘B’, captures the empirical statistics across the 100 samples for the 32 coupling coefficients that connect the 32 primary capsules with the ‘B’ type capsule.

When further matching Panel B with Panel A in Figure 4, one realizes that for each of the six cell types one primary capsule tends to dominate the activation pattern. For example, primary capsule 21 is activated for cell type ‘PDcs’ (both in panel A and B). Similarly, primary capsule 9 is activated for cell type ‘CD16±mono’, while primary capsule 7 corresponds with type ‘T/Mono doublets’, and so on. Note that while the trained model is identical for both Panel A and B, the test sets differ in terms of the input: in Panel A the test set is constructed using all input genes (that is the regular scenario), while for Panel B only the marker genes are used. The latter case is motivated by the idea to have an unobstructed view on how marker genes contribute to primary capsules. This is why the results displayed in Panel B suggest that for each of the six cell types with actionable marker genes, there is one primary capsule that clearly corresponds to these marker genes.

### Cell specific marker capsules

We further performed statistical tests to corroborate the existence of primary capsules that correspond to the six cell types for which marker genes were available. For this, let *t* specify one of the six cell types with marker genes. We further define *m*_*t*_ to be the median of all (32 ×100) coupling coefficients that correspond to type *t*. Let *pc* denote a particular primary capsule. For each such primary capsule *pc*, we construct a 2 ×2 contingency table with values *a, b, c* and *d*, which are defined as follows:

*a* = number of coupling coefficients of primary capsule *pc* having values greater than *m*_*t*_

*b* = number of coupling coefficients of primary capsule *pc* having values smaller than *m*_*t*_

*c* = number of coupling coefficients of primary capsules other than *pc* having values greater than *m*_*t*_

*d* = number of coupling coefficients of primary capsules other than *pc* having values smaller than *m*_*t*_

We then carry out a one-sided (alternative hypothesis = ‘greater’) Fisher’s exact test on the resulting 2×2 contingency table (where a, b, c, d correspond to upper left, upper right, lower left, lower right). Panel C in Figure 4 shows the respective p-values, which have been Bonferroni corrected for multiple testing, for each of the primary capsules *pc* corresponding to the six cell types *t*. Panel C in Figure 4 demonstrates that certain primary capsules are significantly associated with the marker gene expression of the six cells. For example, cell type ‘CD16+ Mono’ is significantly associated with primary capsules 9 and 28, which justifies to annotate them as “marker capsules” because of their responsibility to extract low-level features from the expression of marker genes. The patterns of activation of the primary capsules differ in a way that lets one distinguish the 6 different cell types, such that it is reasonable to use these “marker capsules” as transferring signals for the presence of a particular cell type in a cell population.

Considering the seven cells for which applicable marker genes were not available, primary capsules whose coupling coefficients are significantly increased for a type, may act as “marker capsules” for these types, because the activation patterns of these “marker capsules” allow to clearly distinguish between the different types in the population. As an example, consider types ‘Eryth’ and ‘Memory CD4T’ cells, for which primary capsule 15 and 4, respectively, are activated (see Panel A in Figure 4). This points out that the genes that contribute to primary capsules 15 and 4, and their activation pattern captured by capsules 4 and 15 may serve as marker gene constellations in other applications as well.

In summary, the architecture of the Capsule Networks that we have implemented supports the tracing of genes and their activation patterns that are characteristic for particular cell types. In other words, MarkerCapsule supports the identification of marker genes.

## Discussion

Single cell sequencing technologies have meant major progress in the analysis of the diversity, heterogeneity and variability of cells. Routine exploitation of the corresponding experimental opportunities, however, requires classification systems that enable to determine cell types, states and trajectories of transition between them. Global initiatives (such as the human cell atlas, among in the meantime many others) give evidence of various related activities in research.

Recently, clustering algorithms that compute homogeneous subgroups to potentially reflect identical types/states have been playing a major role. Despite the early successes, clustering entails some substantial drawbacks, because manual annotation is required to assign biological meaning to the groups computed. Therefore, the assignment of meaning relies on existing human knowledge, or at least sufficiently skilled human intuition, to eventually identify types, states and transitions that characterize the cells.

This comes with several issues: first, human knowledge and intuition reflect findings that were predominantly obtained prior to the era of single cell analysis. Hence they cannot take into account the full range of advantages of single cell analysis. Second, because of single cell analysis, the variety of types and states is steadily growing. Therefore, manual execution of the assignment step may require an amount of manpower that is no longer affordable in the meantime.

The integration of heterogeneous data that stem from two (or generally several) simultaneous measurements on single cells reflects most recent experimental progress. This brings up additional methodological challenges: first, unifying data representations are required that enable to combine measurements from two (or more) experimental sources. And, as before, a biologically meaningful interpretation of single cells in the light of such complex data representations is desirable.

Methods that enable to automatically generate useful data representations from complementary measurements on the one hand, and that support automated annotation with types and states on the other hand, are the obvious solution. Methods that additionally allow to track how individual genes or other epi-/genetic effects contribute to the formation of cell types and states are particularly beneficial, because they provide an extra level of biological insight.

Here, we presented MarkerCapsule, designed to address these points and provide novel, adequate and advanced solutions. MarkerCapsule is based on Capsule Networks, which reflect most recent and most advanced progress in deep learning. As such, Capsule Networks first provide full support of automated annotation, by virtue of establishing a supervised learning (in contrast to unsupervised clustering) approach. In that, they do not establish any such approach, but arguably reflect the greatest recent progress in deep learning. Second, although the intended primary purpose was high-performance classification (despite disturbingly blurred or interfering data), Capsule Networks also allowed for human-mind-friendly interpretation of how the input contributed to classification in their original application. Last but not least, Capsule Networks required substantially less data than other methods for training how to annotate unlabeled data.

Our experiments have demonstrated that we were able to re-establish these major, characteristic advances of Capsule Networks also in single cell typing. We have shown that MarkerCapsule outperforms the state of the art approaches in all of these categories: accuracy in basic single cell typing, the amount of annotated data required, and systematic identification of biological meaning that helps to interpret the predictions.

As for the second point, MarkerCapsule evidently requires only a third of the data to achieve utmost accuracy in comparison with other methods. As for the last point, MarkerCapsule has been the first method to relate known marker genes with the corresponding cell types. MarkerCapsule does so in a way that draws an immediate connection with fundamental units of the model, and therefore allows for easy tracking.

In a bit more detail, the first layer of capsules in a capsule neural network (usually referred to as “primary capsules”) is the first layer of units that summarizes the input data in terms of “capsule syntax”. While “capsule syntax” was originally intended to decouple the data from their relative orientation in feature space, which further helped to deal with blurred or mutually interfering data elements, the primary capsules—somewhat surprisingly—proved to convey intuitive meaning as well. Here, we have demonstrated that also in single cell typing, the primary capsules convey a human-mind-friendly, biological interpretation. Namely, primary capsules gathered marker genes whenever such genes existed, which explains the name of the method presented.

In addition to these challenges, MarkerCapsule was able to address how to integrate data stemming from simultaneous experiments on single cells, thereby addressing another important and urgent issue. For that purpose, we demonstrated that variational autoencoder and non-negative matrix factorization based approaches proved to be beneficial when feeding the resulting data representations as input into the underlying capsule networks.

In experiments referring to four different integrated data sets, we evaluated how MarkerCapsule performed in comparison with other methods—note that the other methods were able to process integrated data only because our new methodology also provided suitable data representations for them. MarkerCapsule outperformed the other methods on all data sets. Economy in terms of quantities of data required, and (much enhanced) stability with respect to training data variations were the prevailing advantages. Of note, exactly these advantages meant landmark arguments for capsule networks also in their original application.

In summary, we provided a new method that implemented the latest advances in machine learning (deep learning) for the purposes of typing single cells, both on basic RNA and on integrated, heterogeneous single cell sequencing data. We have demonstrated that the theoretical promises can indeed be leveraged. In this, we argue to have pushed the limits in single cell typing by a non-negligible amount.

We conclude with acknowledging that also our method, of course, leaves room for improvement: various open problems are still awaiting their solution. An important such challenge is the fact that our method, by virtue of being a supervised approach, requires to be provided with annotations prior to classification. While the range of annotations using an automated approach (which we provide) is much enhanced over approaches that require manual intervention at some point, still actionable annotations need to be provided *prior* to running the method.

Methods that are able to not only automatically type, but also automatically derive possible types from the single cell experimental data are likely the next methodological step to be made. For the time being, however, our method may mean an interesting step up in the classification of single cells. In particular, we are first to provide a protocol according to which one can track and understand how input genes (and other genetic effects) combine to cell type defining signals.

## Materials and Methods

### Data Sets

For our experiments, we used four different “multi-omics” single-cell data sets, combining single cell RNA-seq (scRNA-seq) data with other types of single cell sequencing data sets, or complementary scRNA-seq data. The following subsections provide descriptions and links for these four data sets.

### *Data Set 1: CITE-Seq*^*14*^

CITE-seq technology can simultaneously measure different molecular modalities in the same cell. In^14^ CITE-seq is proposed to measure both cellular protein and mRNA expression in one cell, by using oligonucleotide-labeled antibodies. Protein expression and transcriptome profiles establish two distinct and complementary measurements for an individual cell, which justifies to consider the combination of measurements in our experiments. The data set consists of the expression levels of 2000 mRNAs and 13 protein, individually measured in 8,000 cord blood mononuclear cells (CBMCs). We integrated CITE-seq single-cell transcriptomic and proteomic datasets using the autoencoder based approach introduced in the paper^25^, and utilized Seurat^30^, an R package designed for quality control, analysis, and exploration of single-cell genomics data for a preliminary analysis and preprocessing. The preprocessed data contains the expression levels of 2000 genes and ten proteins (3 were removed due to poor quality enrichment) for a total of 7895 cells. As for a specific autoencoder, we utilized the Variational Autoencoder with Concatenated Inputs (CNC-VAE) Architecture^25^. The two datasets have varying dimensions, so we first used two separate encoders and then concatenated the outputs, which was subsequently passed through CNC-VAE to generate the bottleneck layer.

### Data Set 2: scRNA-seq and scATAC-seq from PBMC

Data set 2 provides scRNA-seq measurements (capturing 33538 transcripts) and scATAC-seq measurements in peripheral blood mononuclear cells (PBMCs). We downloaded the corresponding data from 10X genomics and processed the different measurements into a combined representation using Liger^23^, reflecting an NMF based approach, as discussed above. After integration, the data contained 11769 samples (single cells), for each of which the combined representation consisted of 64 features.

### Data Set 3: scRNA-seq from 10X PBMC and scRNA-seq from seq-well

Data set 3 refers to PBMC data sets provided by 10X on the one hand and Giehran et al.^31^ on the other hand, reflecting the sequencing of single cell RNA on two different platforms: 10X and seq-well. Again, the data set is processed using Liger in^23^, resulting in 6713 integrated data samples (single cells) each of which represented in terms of 64 features, as computed by Liger.

### Data Set 4: Mouse brain scRNA-seq and DNA methylation data

The gene expression and methylation data refers to mouse frontal cortical neuron cells. While the expression data refers to 55,803 cells^32^, genome-wide DNA methylation data refers to 3,378 cells^33^. We downloaded the integrated and processed data as originally published^23^, providing *cells*×*genes* count matrices for caudal ganglionic eminence (CGE) interneuron cells.

### State-of-the-Art Approaches: Parameters and Settings

For scPred, we use the R package as provided in the original paper^12^. We use function ‘eigenDecompose’ and ‘getFeatureSpace’ with default parameters (#PC n=10) on the integrated data and use the transformed data as input for the ‘trainModel’ function. We use the ‘scPredict’ function of the scPred framework on test data with a threshold 0.7 (we chose 0.7 over the default 0.9, because we found 0.7 to give optimal results for scPred, results for the default 0.9 were rather poor in particular). For comparing accuracies, we treat the ‘unassigned’ label as an incorrect label, as mentioned in the original paper of scPred^12^.

For AcTINN, we use the Python package provided in the corresponding Github repository. We apply default hyperparameters as provided in the package for training the model.

For CHETAH, we use the bioconductor package ‘CHETAH’ available at the corresponding Github repository. We use training data as a reference dataset and test data as input in the ‘CHETAHclassifier’ function.

### Capsule Networks: Foundations and Architectural Elements

Capsule networks can be interpreted as neural networks that process vectors instead of scalars, where capsules, as their defining entities, reflect arrangements of scalars (ordinary neurons) into vectors (capsules). The length of the capsule vector reflects the probability that the entity to which the capsule refers is present in the input object. In the basic version of Capsule Networks, two layers of capsules are used: lower-level capsules (“primary capsules”) reflecting the most basic entities of an object are connected to higher-level capsules (“digit capsules” in the original application; see below) that combine the input of the lower-level (primary) capsules into information that enables to classify the object. This mirrors the situation that early layers of neurons provide their output as input to subsequent layers in standard neural networks.

In the original application, the recognition of digits in images, each higher-level capsule referred to one of the ten different digits from 0 to 9, and capsule networks outperformed conventional CNN’s greatly when digits overlapped/shaded each other or were distorted otherwise. The length of each of the capsule vectors reflected the probability that the digit was present in the image. Lower-level capsules (“primary capsules”) that provided their output as input to digit capsules coded for orientation, width, scale, thickness, etc. (where ordinary neurons can only code for intensity and color). Beyond the enhancement of classification itself, the human intuition friendly interpretation of lower-level capsules was perceived as a particular advantage of CapsNets.

In our application, higher-level capsules immediately correspond to cell types. As pointed out in Results, we found primary capsules to reflect marker genes for types for which such marker genes were known. This means that also in single cell type classification, primary capsules enjoy a human mind friendly interpretation. More than that, they may suggest plausible sets of marker genes for cell types for which markers had not been available so far.

As an important theoretical fact, the *viewpoint invariance property* of CapsNets ensures that orientation and relative position of objects and entities can be captured by virtue of the design of CapsNets. While this property is favorable for various reasons, a consequence of particular importance is that they require substantially less training data than other approaches. The explanation is that alternative approaches often need to be presented with training data viewed from various angles, while CapsNets only need to be presented with one datum during training. This explains why CapsNets require substantially less training data for achieving superior performance^24^, which our Results testify as well.

### Dynamic Routing

Providing the output of earlier layers of CapsNets, consisting of primary capsules, to later layers, consisting of “cell type capsules” is implemented by a *dynamic routing* protocol. The protocol can be interpreted as determining for a primary capsule which higher level capsule optimally accommodates the output of the primary capsule. According to this, in our application, cell type capsules are activated if primary capsules (shown to correspond to marker genes if such genes exist) support their activation. In other words, in MarkerCapsule, cell type capsules are activated, if the corresponding marker capsules become active. In other words, a cell type is recognized if the gene expression values include gene activation patterns that are characteristic for the type.

### MarkerCapsule Network Architecture: Details

The architecture of the proposed model is shown in Figure 6. The model consists of two convolutional layers: Conv1, and PrimaryCaps and one fully connected ‘cell-type’ capsule layer. The first convolution layer converts the (possibly integrated) input data for each sample into the intermediate-level feature representation, which in turn serves as input for the primary capsule layers and subsequently the ‘cell-type’ capsule layer for further feature abstraction.

**Figure 6.**
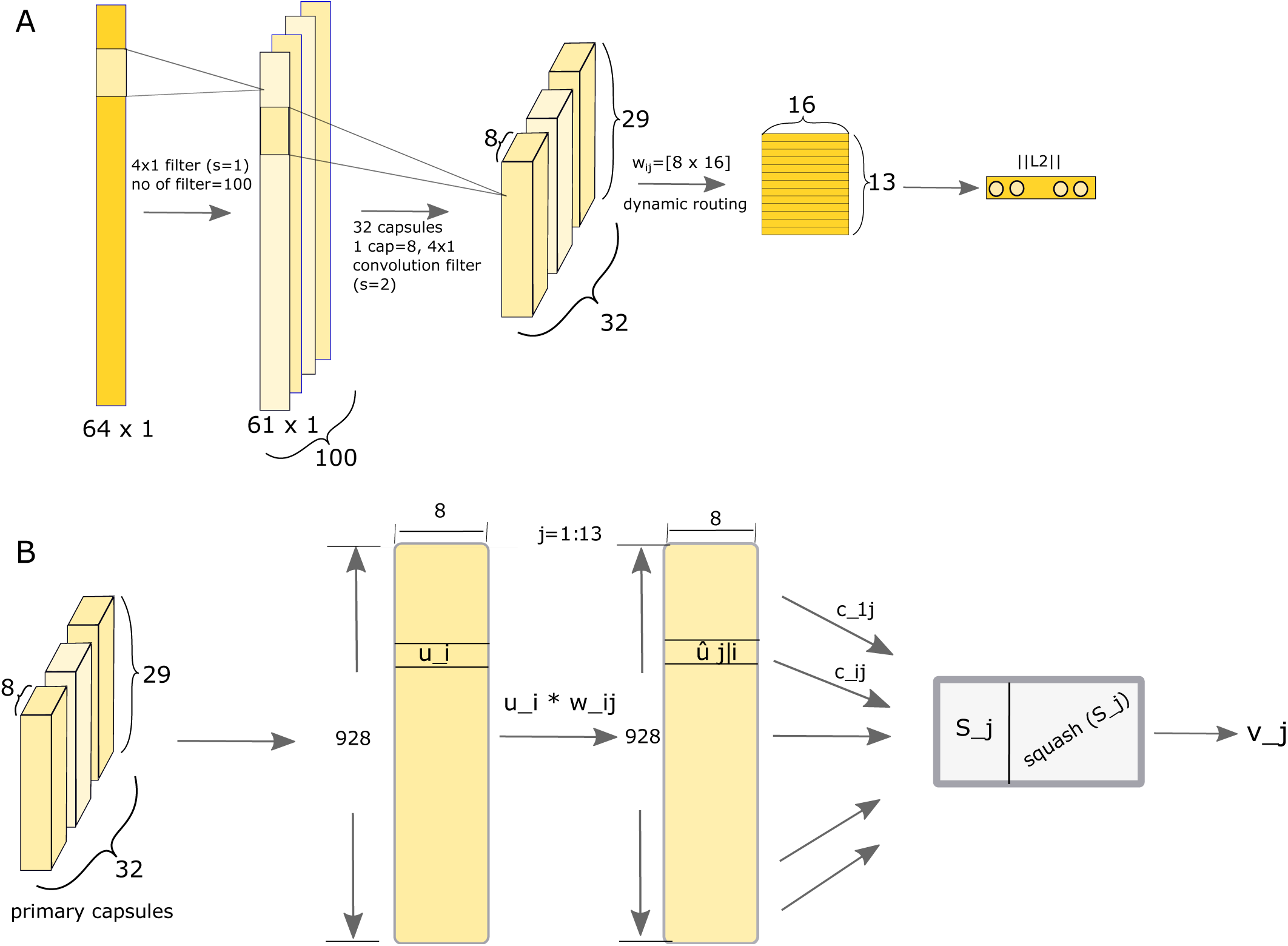
Figure shows the details of different layers of the proposed capsule model architecture. The first layer is a convolution layer which is followed by 32 primary capsules, each having 84 ×1 convolution filter. The higher layer consists of 13, 16D ‘cell-type’ capsules representing the 13 unique cell types of the dataset CITE-seq. The L2-norm of each capsule is calculated which represents probability of the presence of that particular cell type. Panel-B shows the internal computation between primary capsules and ‘cell-type’ capsules. Primary capsule layer can be viewd as there are 928 capsules(*u*_*i*_) of 8 dimension. For a particular higher level capsule *j* each *u*_*i*_ is multiplied with weight *w*_*i j*_ of dimension 8 ×10 to produce *û*_*j*_|_*i*_. The output *v* _*j*_ is produced by a weighted sum over all *û*_*j*_|_*i*_ followed by squashing nonolinear activation function. The coupling coefficients *c*_*i j*_ are determined by the dynamic routing algorithm.

The first convolution layer consists of 100 convolution filters of size 4×1 applied at a stride of 1 (at no padding), followed by ReLU activation^34^, which processes the (pre-processed) input data of size 64×1. Because no padding is applied the input data is turned into a 100×61×1-tensor.

As described in the seminal paper^24^, a primary capsule also incorporates convolutional principles. Here, the primary capsule layer consists of 32 capsules each of which incorporates eight 4×1 filter operating at a stride of 2, taking in the 100×61 ×1-tensor, and generating output consisting of 32 units, each of which refers to one primary capsule and reflects a 29×1 ×8-tensor, as follows from the principles of capsules (the dimensionality of 8, reflecting vectors of length 8, is adopted from the seminal paper^24^).

The purpose of primary capsules is to extract the basic lower level features from the input of the convolution layer. The length of the 8-dimensional vectors emerging from the primary capsules reflects how likely the entity is present in the data, and as usual^24^, we apply a “squashing” activation function that scales the length of the vector into the range [0, 1], so as to put the lengths of the vectors into a one-to-one context with probabilities. In some more detail, the squashing function applied here performs that scaling 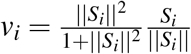 where *v*_*i*_ is the vector output of capsule *i*, and *S* _*i*_ is its input.

Squashing activation is also applied in the higher-level layer of ‘cell-type’ capsules. We have 13 higher-level capsules or ‘cell-type’ capsules, each of which has 16 dimensions. For ‘CITE-seq’ data, we have 13 unique types of cells, and for each type, we have one ‘cell-type’ capsule. For other datasets we keep the same architecture of the model except the number of higher level capsules. The length of the vector output of the ‘cell-type’ capsule represents the probability that a cell is of that particular type. The cell type capsules receive the input from all primary capsules, and by applying the dynamic routing protocol, the coupling coefficients between the primary capsules and the output capsules are determined. As each primary capsule outputs a 29×1×8-tensor, and there are 32 capsules, we can view this as 29 ×32 = 928 8-dimensional (8-D) vectors, which serve as input for the ‘cell-type’ capsules.

Panel B in Figure 6 sketches how the data is transformed when passed on from the primary to the cell type capsules. Here we define *u*_*i*_, *i∈* [1, 928] as the 8-D capsule in the primary capsule layer, *w*_*i j*_ is the weight matrix of dimension 8×16 that performs the affine transformation of the input from the primary capsule layer. The input of the ‘cell-type’ capsule *j* where *j∈* [1, 13] is the affine transformed vector *û*_*j*|*i*_ = *u*_*i*_*\* w*_*i j*_ where *I ∈* [1, 928]. The dynamic routing protocols return vector *v* _*j*_ for each of the ‘cell-type’ capsule *j*, which represents the probability of the existence of a particular cell type *j*. In Panel B of Figure 6, we can see that *v* _*j*_ is produced by a weighted sum (*s* _*j*_) over all the *u* _*j*_|_*i*_ as 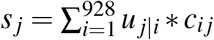, where *c*_*ij*_ signifies the coupling coefficient between primary capsules (*i*) and ‘cell-type’ capsules (*j*), which are determined by the dynamic routing protocol^24^.

The weighted sum is subsequently transformed into the range [0, 1] using the squashing activation function, as described above. The coupling coefficient *c*_*i j*_ are to sum to 1 across all primary capsules *i*. For example, for cell type capsule *j*, 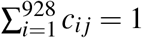. The updating algorithm of *c*_*i j*_ is primarily governed by a “sense of agreement” between *v* _*j*_ with *u* _*j*|*i*_ and is implemented as a dot product between them. The primary capsule *i* sends its output (*u* _*j*|*i*_) to that higher level capsule *j*, which agrees with its output (*v* _*j*_)· *c*_*i j*_ is updated by the formula *c*_*i j*_← *c*_*i j*_ + *u* _*j*|*i*_*· v* _*j*_. Computing the dot product between *u* _*j*|*i*_ and *v* _*j*_ determines whether *c*_*i j*_ increases or decreases, depending on the angle between *u* _*j*_|_*i*_ and *v* _*j*_. In summary, the coupling coefficient *c*_*i j*_ can be interpreted as the likelihood of cell type *j* applying given the lower level features extracted/captured by primary capsule *i*.

## Notes

### Competing Interest Statement

The authors have declared no competing interest.

